# Feedback ratiometric control of two microbial populations in a single chemostat

**DOI:** 10.1101/2021.03.05.434159

**Authors:** Davide Fiore, Fabio Della Rossa, Agostino Guarino, Mario di Bernardo

## Abstract

We address the problem of controlling the dilution rate in a chemostat to regulate the ratio between the concentrations of two microbial populations growing in continuous culture. After analyzing the open-loop dynamics of this multicellular system, we present two alternative feedback control strategies, one based on a gain-scheduled state feedback controller, the other on a switching control strategy. We show that both strategies are effective in solving the problem and illustrate the results by a set of representative numerical simulation.

## I. Introduction

Synthetic Biology aims at engineering biological systems with new functionalities, with applications ranging from health treatments to bioremediation, to production of biofuels and drugs in bioreactors. This is made possible by embedding into living cells, such as bacteria and fungi, artificial genetic circuits modifying the natural behavior of these biological systems. That is, they change at what condition and rate genes are expressed in the cells to produce proteins or chemicals. However, the complexity and hence the capabilities of these engineered genetic circuits are limited by inherent factors in their host cells, such as excessive metabolic burden, competition of limited resources and incompatible chemical reactions. To overcome these limitations a promising strategy is to distribute the required functionalities among several cell populations forming a microbial consortium, so that each cell strain embeds a smaller subset of engineered genes [1]–[4]. In this way each cell carries out a specialized role and, by dividing the labor, contributes more efficiently to the achievement of the final goal. Unfortunately, this solution comes with an additional challenge; due to unavoidable differences in their genetic loads, each cell population in the consortium will grow at different rates, and can give rise to undesired dynamics, such as oscillations, or even extinction [5]. Therefore, for the correct operation of the designed consortium it is essential to guarantee stable coexistence between the constituent populations by regulating their relative numbers. Moreover, in applications in which each population produces a different chemical reagent, keeping the balance in the consortium at the correct ratio is fundamental to achieve high efficiency in the production of desired metabolic end products. In the absence of mechanisms designed *ad hoc* into the cells allowing them to self-balance their ratio, it is necessary to regulate the populations’ densities by means of some external action [6]. Also, in industrial applications where high production efficiency is required, external control strategies should be preferred to embedded solutions, because additional genetic circuits can cause further metabolic burden to the cells and hence lower production rates of the desired metabolic end products.

Previous work presented in the literature considered the problem of guaranteeing coexistence of two cellular populations in a chemostat. In [7] the authors showed that two species can be made to coexist by means of state-feedback control of the dilution rate, with the resulting closed-loop system having an equilibrium point in the nonnegative orthant to which all solutions converge. The proposed controller was shown to be robust with respect to bounded uncertainties on the growth functions, however the closed-loop dynamics could not be arbitrarily assigned due to constraints on the control gains. Moreover, the problem of guaranteeing survival of a resident species with respect to the invasion of a new species was also studied in [8].

In this letter we consider the problem of regulating the relative numbers, i.e., the *ratio*, of two independent cellular species in a chemostat, while, at the same time, guaranteeing their survival and the convergence dynamics of the closed-loop system. After presenting a complete description of the rich nonlinear dynamics of the system, we propose and analyze two control algorithms that, by acting only on the dilution rate, can regulate the ratio between the concentrations of the two species grown in continuous culture in the same chemostat to a given desired value, say *r*_d_. The first control algorithm we present is a gain-scheduled state-feedback controller guaranteeing global convergence to the desired set-point. The second control algorithm is a switching controller changing the dilution rate between two values to guarantee convergence of all solutions towards the desired set-point located on the sliding region. We analyze the closed-loop dynamics for both controllers and compare their robustness to parameter perturbations. Finally, we conclude by discussing the advantages and limitations of the proposed control strategies.

## II. Mathematical models and Dynamics

We consider two microbial populations growing in continuous culture in a chemostat [7], [9]. The two species are assumed to feed on the same substrate and to be independent, that is, they do not directly influence their respective growth rate. The chemostat model with two species, can be expressed as

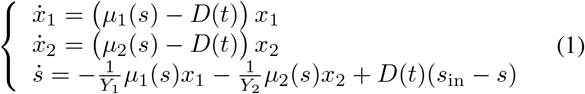

where the variables *x*_1_, *x*_2_ ∈ ℝ_≥0_ denote the concentrations of biomass of the species 1 and 2, respectively, and *s* ∈ [0, *s*_in_] denotes the concentration of the substrate in the chemostat. Moreover, *μ_i_*(*s*) is the growth rate of the *i*-th species, *s*_in_ is the concentration of substrate in the inlet flow, assumed to be constant, and *Y_i_* is the yield coefficient, defined as the ratio between the biomass created and the substrate consumed, assumed without loss of generality to be unitary for both species. The control input *D*(*t*) : ℝ_≥0_ ↦ [*D*_min_, *D*_max_], with *D*_min_ > 0, is the dilution rate, defined as the ratio between the inlet flow rate and the culture volume in the chemostat. The dilution rate is assumed to be the same for both species, that is, the culture is well-mixed and mortality and attachment of the bacteria are neglected [8].

Under the assumption that the control input *D*(*t*) is persistently exciting [10], i.e., such that 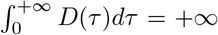, it follows that the subspace 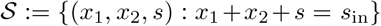 is attractive and invariant for all solutions of system (1). So, provided that *D*(*t*) > 0 for all *t* > 0 and that the initial condition of system (1) belongs to 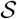, the dynamics of (1) on 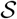 can be described by considering the *reduced system* [7]

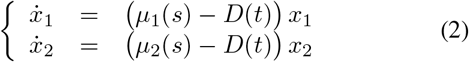

defined on the invariant set 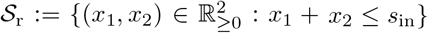 (Fig. 1).

**Fig. 1.**
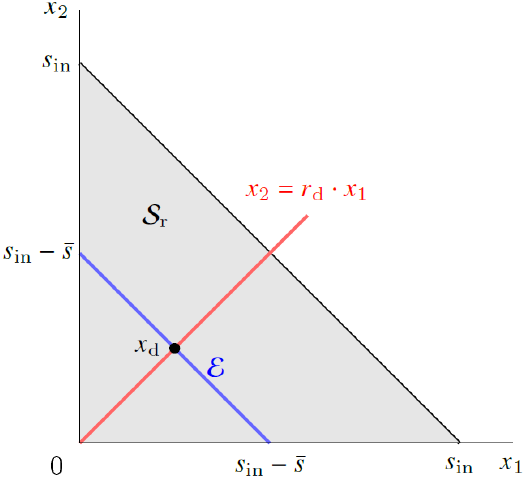
Graphical representation of the domain of definition 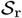 (gray region) of the reduced system (2) and position of the control set-point *x*_d_. The set-point *x*_d_ is the point of intersection between the equilibrium set 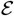 defined in (6) (blue line) and the line designing the desired ratio *r*_d_ defined in (7) (red line).

### A. Growth dynamics

A fundamental role in the dynamics of model (1) is played by the growth rate functions *μ_i_*(·) : [0, *s*_in_] ↦ ℝ_≥0_, which are generally assumed to be *C*^1^, strictly increasing and such that *μ_i_*(0) = 0, for *i* = 1, 2. We consider the case in which *μ*_1_ and *μ*_2_ have at most two intersection points, namely at *s* = 0 and 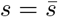, that is the case of strictly concave growth rate functions. When they intersect only at *s* = 0, it follows that one species grows always faster than the other, that is, *μ_i_*(*s*) > *μ_j_*(*s*), *i* ≠ *j*, for all *s* > 0 and for any value of *D*. From the Competitive Exclusion Principle (CEP) [11], this implies that no control input *D*(*t*) exists such that more than one strain survives at steady state. Specifically, any positive solution of (1) converges to the equilibrium point such that *s* = λ_*i*_(*D*), *x_i_* = *s*_in_ – λ_*i*_(*D*) and *x_j_* = 0, where λ_*i*_(*D*) := sup{*s* ∈ [0, *s*_in_] : *μ_i_*(*s*) < *D*}, *i* = 1, 2, is the so-called break-even concentration.

On the other hand, if the growth functions also intersect at 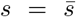, it is possible for the two species to coexist. This corresponds to a scenario in which the two species are complementary, that is, one species grows faster than the other for low values of *s*, while the situation is reversed for high values of s. In what follows we will consider this latter scenario obtained by making the following assumption [8].

#### Assumption 1

There exists 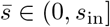 such that

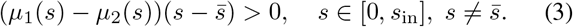

Moreover, we denote with 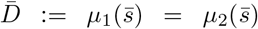 the corresponding value of the dilution rate.

In the rest of this letter we consider growth functions of Monod’s type [12], that is, we assume

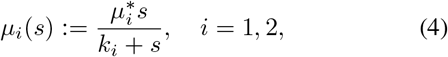

where 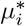 and *k_i_* are positive constants.

### B. Open-loop dynamics and bifurcation analysis

Before presenting the control problem and the solutions we propose in the next sections, we analyze here the rich nonlinear dynamics of the open-loop system (2). Namely, we show the steady state behavior of system (2) and how it changes as the dilution rate *D* is being varied.

The equilibrium points of system (2) can be obtained as the intersections between the four nullclines of its vector field in the domain 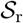. Specifically, each equation 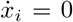, *i* = 1, 2, has two solutions, namely *x_i_* = 0, that is, the *x_j_*-axis, *i* ≠ *j*, and

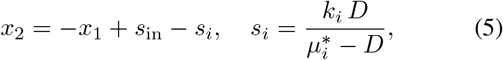

where the last equation follows from *μ_i_* being of Monod’s type (4). Therefore, the origin (0, 0) is always an equilibrium point for the system; its stability depending on *D*. Moreover, the system might admit other steady states whose number and stability depend on the values of *D* and on the growth functions *μ_i_*. Specifically, we can identify the following six cases corresponding to different positions of the two nullclines (5) in 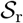 (Figure 2):

I. *D* = 0: when *s*_1_ = *s*_2_ = 0 the nullclines (5) overlap and all solutions asymptotically converge to the stable equilibrium set 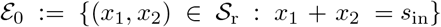, while the origin is locally unstable. This case corresponds to an undesired working condition in which no new substrate is added to the reactor and the biomass is in starvation. However, this situation can never occur in continuous culture as it is always assumed that *D*(*t*) > 0, ∀*t* > 0.
II. 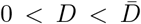: there are two equilibrium points, each one on an axis, corresponding to their intersection with the nullclines (5). Specifically, 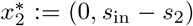 is a stable node and 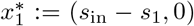 is a saddle, where sj is as in (5), while the origin is locally unstable. This case corresponds to a low concentration of the substrate at steady state due to its consumption and low dilution rate. This results in species 2 prevailing over species 1.
III. 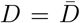: the nullclines (5) again overlap and all solutions asymptotically converge to the stable equilibrium set

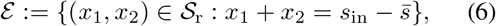

while the origin is locally unstable. These equilibrium points correspond to a condition of stable coexistence between the two species.
IV. 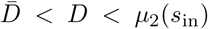: this case is similar to case (II) but with opposite stability properties of the equilibrium points, that is, 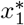 is a stable node and 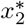 is a saddle, while the origin is still locally unstable. In this case species 1 prevails at steady state because of the high concentration of substrate.
V. *μ*_2_(*s*_in_) < *D* < *μ_i_*(*s*_in_): there is only one intersection in the domain 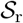 between the nullclines (5) and the axes at the point 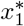, which is stable, while in this case the origin is a saddle. At steady state species 2 is washed-out from the chemostat due to the dilution rate *D* being greater than its maximum growth rate *μ*_2_(*s*_in_).
VI. *μ*_1_(*s*_in_) < *D* < *D*_max_: there is a unique stable equilibrium point in the origin to which all solutions converge. In this case, *D* being greater than the maximum value of both growth rates causes the washout of both species, that is, the cells are removed from the chemostat faster than they can grow.

**Fig. 2.**
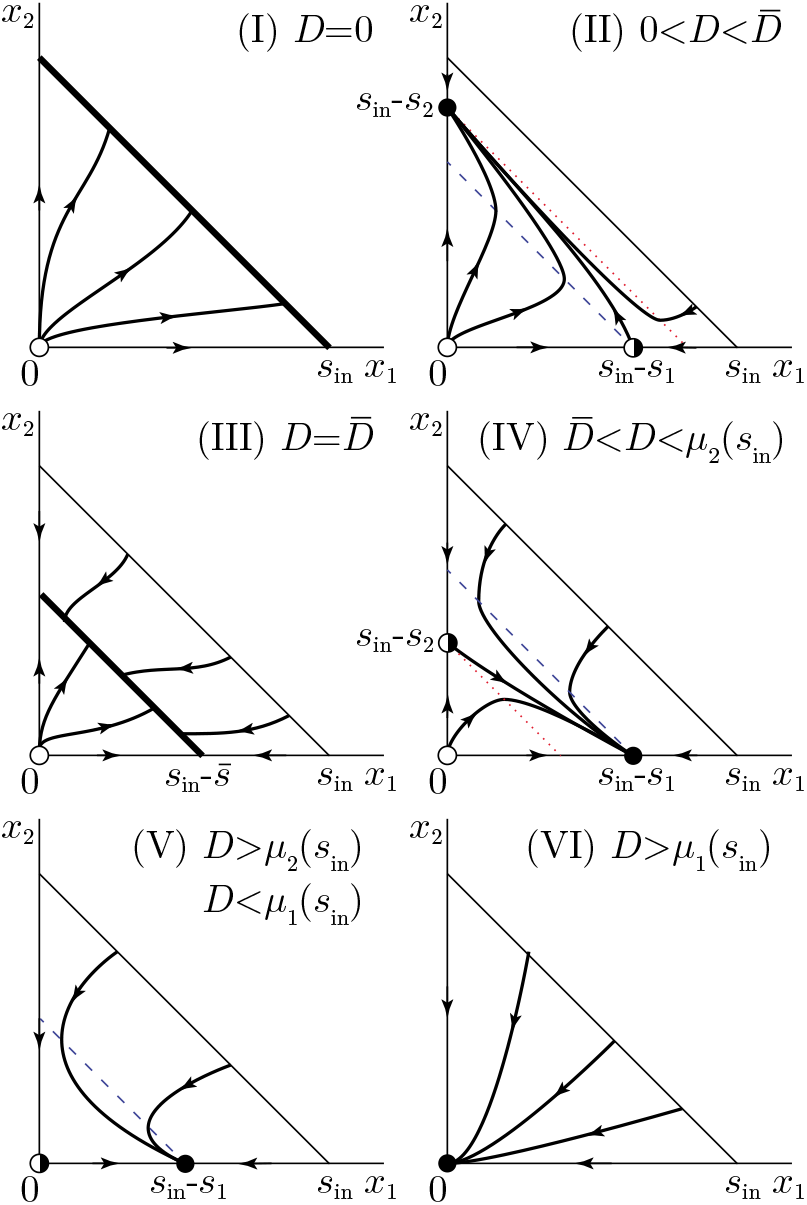
Possible state portraits of system (2) for different values of constant dilution rate *D*. Blue dashed and red dotted lines correspond to the two nullclines of the system reported in (5) for *i* = 1 and *i* = 2, respectively. Full, empty, and half-filled dots represent stable, unstable and saddle equilibria, respectively.

The transitions between the dynamical behaviors described above are due to bifurcations of the equilibrium points of the system. In particular, in cases (I) and (III) the system undergoes a *degenerate transcritical bifurcation* [13], in which the equilibrium sets 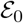 and 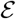 appear, respectively, as the two nullclines (5) overlap. These two sets are not structurally stable, since they suddenly disappear for any small perturbation of the bifurcation parameter *D* from the bifurcation points *D* = 0 and 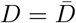. On the other hand, for *D* = *μ*_2_(*s*_in_) (*D* = *μ*_1_(*s*_in_)) the equilibrium points undergo a regular transcritical bifurcation, in which the equilibrium point 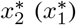 collides with the one in the origin, exchanges stability and exits the domain 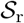.

## III. Control Problem formulation

Under Assumption 1 it follows that, by applying a constant, open-loop control input 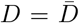 to system (2), coexistence of the two species can be achieved at steady state. Specifically, any solution of (2) converges to the equilibrium set 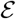 defined in (6), and more precisely to a point whose position depends on the initial condition *x*_0_ = (*x*_0,1_, *x*_0,2_). Therefore, it is not possible by means of any open-loop control input to guarantee at the same time coexistence of both species while regulating the *ratio* between the steadystate concentrations *x*_1_ and *x*_2_ to some reference value. Moreover, structural instability of the equilibrium set 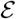, uncertainties on the parameters of the growth functions *μ_i_*(*s*) and other disturbances cannot be rejected by means of any open-loop control law. It is therefore necessary to regulate the dilution rate of the chemostat by employing closed-loop control laws. More formally, we want to solve the following control problem.

### Problem statement

Design some feedback control input *D*(*x*_1_, *x*_2_) for system (2) such that:

1. *coexistence* of the two species is guaranteed for all time, that is, *x_i_*(*t*) > *x*_*i*,min_, ∀*t* > 0, *i* = 1,2, where *x*_*i*,min_ > 0 is some safety threshold to avoid extinction of species *i* due to unwanted events, such as washout or starvation;
2. the *ratio x*_2_/*x*_1_ between the concentrations of the two species is robustly regulated at steady state to a desired value *r*_d_ > 0, that is,

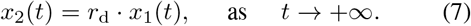

Notice that the previous control problem corresponds to requiring that all solutions of system (2) are stabilized to the point of intersection between the equilibrium set 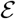 in (6), where coexistence is possible, and the line defined by (7) (see Fig. 1), that is, to the point

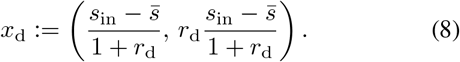

We further assume that the concentrations *x*_1_ and *x*_2_ are either directly or indirectly measurable, for example by means of fluorescent reporter proteins produced by one or both species.

## IV. Proposed controllers

In this section we present and analyze two feedback control strategies that solve the control problem presented in Section III. Namely, a proportional gain-scheduled state-feedback and a switching controller. After presenting the control laws, we validate and compare their robustness by means of numerical simulations.

In all simulations we use the following values of the parameters of the growth functions *μ_i_*(*s*) [8]: 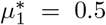, *k*_1_ = 5, 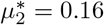, *k*_2_ = 0.13. Furthermore, we set *s*_in_ = 5, *D*_min_ = 0.1, *D*_max_ = 1. These limiting values of the dilution rate *D* guarantee, respectively, a continuous flow rate of culture medium and that fresh medium equal to the volume of the culture is added in the bioreactor at most in one hour.

### A. State-feedback control

The first controller we propose is a state-feedback controller whose control gains continuously depend on the desired ratio *r*_d_. This to guarantee, locally to the set-point *x*_d_, the same closed-loop dynamics for any *r*_d_. Specifically, the control input is defined as

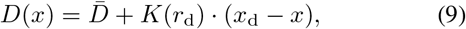

saturated when exiting the interval [*D*_min_, *D*_max_], with *K*(*r*_d_) = (*K*_1_(*r*_d_), *K*_2_(*r*_d_)) being the control gain vector. The control input (9) is composed by two terms: (i) a feedforward action 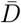, that guarantees convergence to the equilibrium set 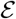 of coexistence between the two species, and (ii) a feedback action *K*(*r*_d_) · (*x*_d_ – *x*) that steers the state *x* towards the desired set-point 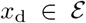 and rejects disturbances due to uncertainties on the system parameters.

The control gain vector *K* (*r*_d_) is evaluated by assigning, via pole placement, the desired eigenvalues (λ_1_, λ_2_) to the closed-loop dynamics (2) and (9) locally to the desired setpoint *x*_d_. Specifically, the control gains are obtained by solving the equations tr(*J*) = λ_1_ + λ_2_ and det(*J*) = λ_1_λ_2_, where *J* is the Jacobian matrix of the linearized system 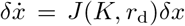, with *δx* = *x* – *x*_d_. They can be given as

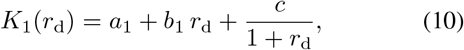

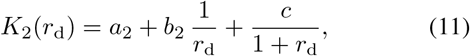

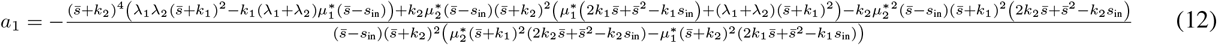

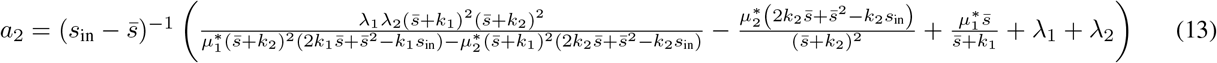

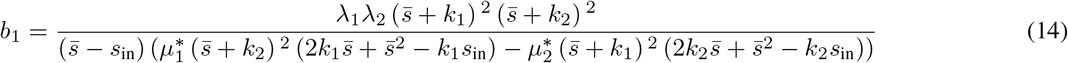

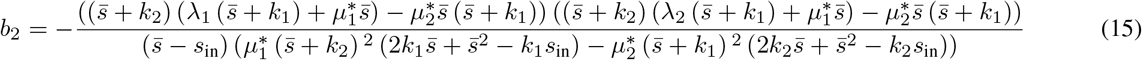

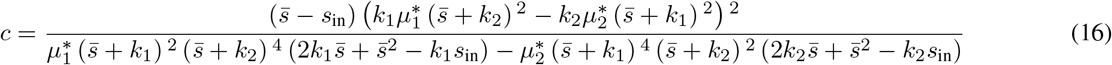

where *a*_1_, *a*_2_, *b*_1_, *b*_2_, *c* are coefficients whose expressions are defined in (12)-(16).

By numerical analysis, the closed-loop dynamics has been found to have four equilibrium points: a stable node in *x*_d_, an unstable node in the origin, and two saddles, each on one different axis (Fig. 3, left panel), and their stability properties do not depend on the particular choice of the control parameters. Therefore, under the action of the feedback control input (9), all solutions converge to *x*_d_ for any initial condition 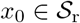.

**Fig. 3.**
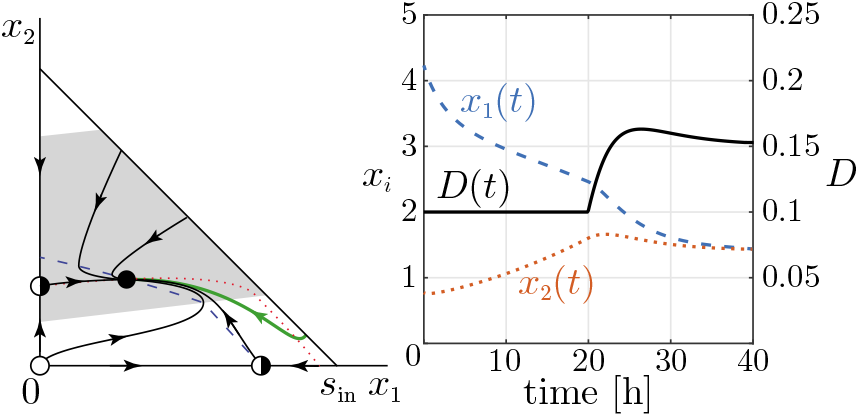
State-feedback controller. Left panel: State portrait of the closed-loop system (2) and (9). Blue dashed and red dotted lines correspond to the two nullclines. Full, empty, and half-filled dots represent stable, unstable and saddle equilibria, respectively. The gray area of the plane represents the (*x*_1_, *x*_2_) values in which the control input is not saturated. Right panel: time evolution of the state and of the control input of the thick green trajectory reported in the left panel. For the closed loop system, λ_1_ = λ_2_ = −0.2, *r*_d_ = 1, *x*_d_ = (1.42, 1.42), *K*_1_ = 0.11, and *K*_2_ = −0.34.

A representative simulation of the closed-loop system is reported in Fig. 3. The control strategy is able to regulate the two concentrations to the desired ratio *r*_d_ = 1, as required; the state of the system converging to the point *x*_d_ lying on 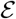. Notice that the response of the closed-loop system is largely affected by the intrinsic growth rate of the biomass in the chamber which, of course, cannot be sped up by the control input.

### B. Switching control

The second controller we consider is a switching controller that changes the control input *D* between two constant values, *d*^+^ and *d*^−^, with *d* < *d*^+^, depending on the sign of a scalar function *σ*(*x*), that is,

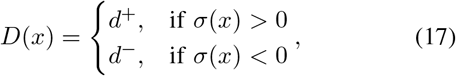

where

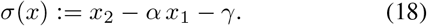

The switching surface Σ is defined as the zero-set of *σ*(*x*), i.e., 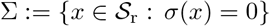, and it is chosen such that Σ and 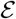 intersect at the point *x*_d_ in (8), linking the parameters *α* and *γ* as follow

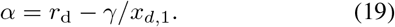

Moreover, if the control parameters are chosen such that

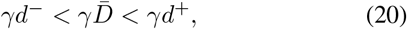

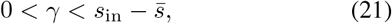

then (i) there exists a sliding region 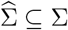 towards which all solutions converge, and (ii) *x*_d_ is a stable pseudo-equilibrium point for the sliding vector field *f_s_*(*x*).

Next we prove the previous statements. We first analyze (i). Denoting with *f*^+^ (*x*) and *f*^−^(*x*) the closed-loop vector fields when *D* = *d*^+^ and *D* = *d*^−^, respectively, and with 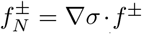 their normal component to Σ, a sliding region exists if 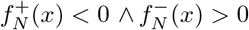 for some *x* ∈ Σ. From the first condition on 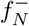 and ∇*σ* = [–*α* 1]^⊤^ we have

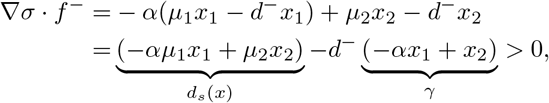

which is equal to *d*^−^*γ* < *d_s_*(*x*). Likewise, from the second condition on *f*^+^ we get *d_s_*(*x*) < *d*^+^ *γ*. Combining these two previous conditions we obtain

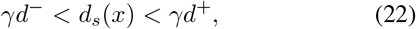

where *d_s_*(*x*) := −*αμ*_1_(*s*)*x*_1_ + *μ*_2_(*s*)*x*_2_. Also, using Filippov’s convention [14, p.52], i.e.,

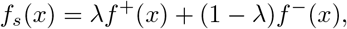

with

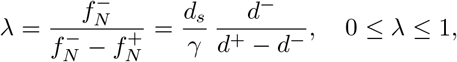

the sliding vector field can be defined as

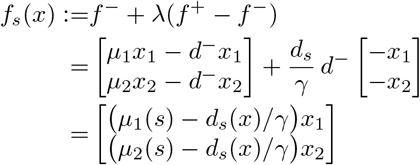

and it has four equilibrium points, namely (0,0), (0, *γ*), (−*γ*/*α*, 0), and alastone for 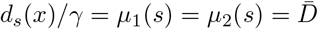 at the point of intersection between Σ and 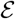. Therefore, using (22) we have that the desired set-point *x*_d_ is a pseudoequilibrium point of *f_s_* only if the control parameters satisfy (20). Moreover, condition (22) also implies that *γ* > 0.

To prove (ii) we can analyze the local stability of the equilibrium points in (0,0) and (0, *γ*) of the sliding vector field *f_s_*(*x*) by studying the sign of its components locally to the points. Indeed, if these two points are found to be unstable, then the pseudo-node in *x*_d_ must be stable. It can be proved that the origin is always unstable since *f*_*s*,1_(*x*_1_, 0) and *f*_*s*,2_ (0, *x*_2_) are both positive close to the origin. For what concerns the point in (0,*γ*), we have that *f*_*s*,2_(0, *x*_2_) > 0 for 0 < *x*_2_ < *γ* and *f*_*s*,2_(0, *x*_2_) < 0 for *x*_2_ > *γ*, therefore solutions on the *x*_2_-axis converge to (0, *γ*). on the other hand, in the transversal direction we have that

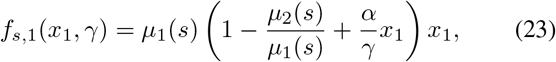

whose changing of sign depends on *γ* being greater or less than 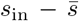, since the term 1 – *μ*_2_(*s*)/*μ*_1_(*s*) in (23) is positive only when the state *x* is below 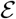, that is, when 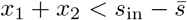. Hence, it follows that when 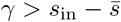 the point (0, *γ*) is a stable node, while when 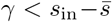 it is a saddle. Therefore, the equilibrium point *x*_d_ at the intersection between Σ and 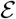, is stable if condition (21) holds.

Note that at the boundary of this region in the parameter space two different bifurcations occur. When 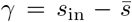 the stability of the desired point is lost due to a degenerate transcritical bifurcation similar to the one described in the previous section. As *γ* decreases, each of the two vector fields generates a *fold bifurcation of tangent points* [13] of the Filippov vector field, which gives rise to crossing regions in the switching surface at the right and at the left of the pseudo-equilibrium, respectively (see the crossing region at the left of the pseudo-equilibrium in the lower-left panel of Fig. 4), causing the boundaries of the sliding region 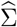 to shrink, until 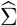 disappears for *γ* = 0 at an *invisible-invisible pseudo-Hopf bifurcation* [13]. Note that if *γ* = 0 the bifurcating pseudo-equilibrium is stable, and is reached via converging crossings of the switching manifold, at a rate less than exponential. However, for *γ* in (21) there always exists an attractive sliding region in the neighborhood of the pseudo-node at *x*_d_, to which all solutions in 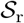 converge.

**Fig. 4.**
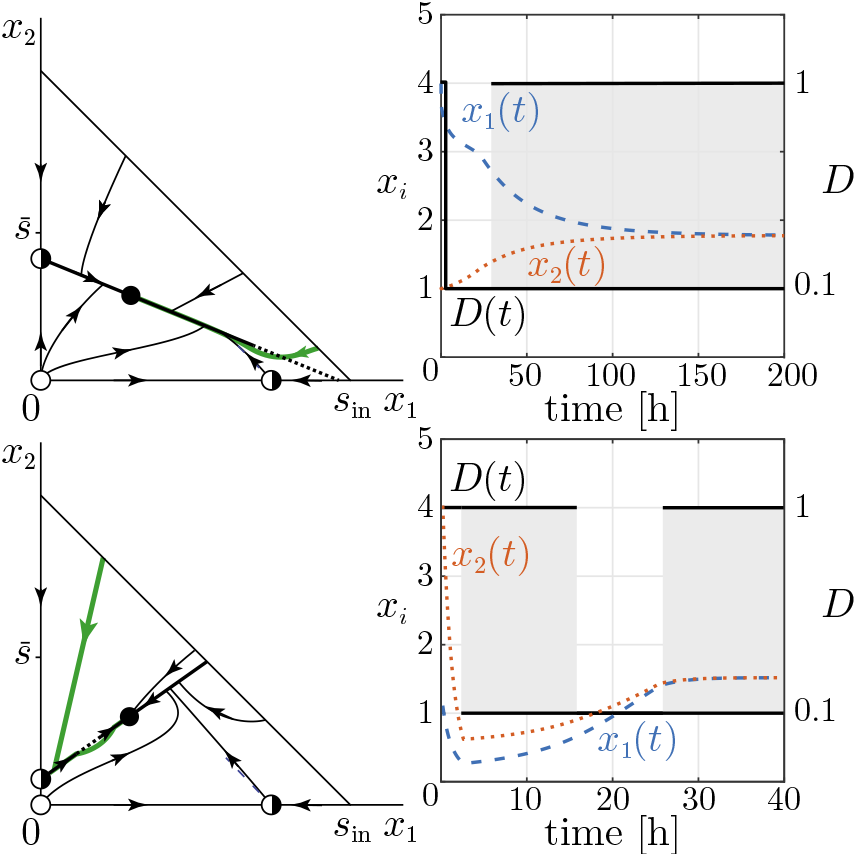
Switching controller. Left panels: State portraits of the closed-loop system (2), (17) and (18). The switching surface Σ is the thick line, full when sliding conditions (22) are satisfied, dotted otherwise (crossing region). Full, empty, and half-filled dots represent stable, unstable and saddle equilibria or pseudo-equilibria, respectively. In the top panel *γ* = 2, in the bottom panel *γ* = 0.4. Right panels: time evolution of the state and of the control input of the thick green trajectories reported in the respective left panel. In the shaded regions, the control switches at high frequency between the two values. Other parameter values: *r*_d_ = 1, *x*_d_ = (1.42, 1.42), *d*^+^ = *D*_max_, *d* = *D*_min_.

Two representative simulations of the closed-loop system are reported in Fig. 4, for choices of *γ* > *x*_*d*,2_ (top panels) and *γ* < *x*_*d*,2_ (bottom panels). In both cases there exists a unique stable pseudo-equilibrium point, on the sliding region 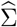 at its intersection with the equilibrium set 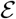. Notice that in the case *γ* > *x*_*d*,2_ the convergence of sliding solutions to the pseudo-equilibrium is slower than in the other case. This is essentially due to *f*^+^(*x*) and *f*^−^(*x*) pointing almost in the same direction in the neighborhood of 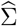, and does not depend significantly on the particular choice of *d*^+^ and *d*^−^ being made. For this reason and to guarantee greater robustness to disturbances, as we are going to show next, the control parameter *γ* must be chosen in the range

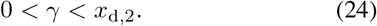

In conclusion, to stabilize all solutions on the set-point *x*_d_, it suffices to choose the control parameters in (17)-(18) so as to satisfy conditions (20), (19) and (24).

### C. Robustness and performance analysis

Here we first analyze the robustness of the proposed controllers to variations of the parameters of the growth functions *μ_i_*(*s*), *i* = 1,2, and later we compare their performance in terms of average settling time. Perturbations on the values of the parameters 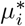 and *k_i_*, can cause the growth functions to intersect each other at a value, say 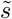, different from the nominal value 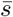. This in turn results in a change of the position of the equilibrium set 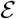 defined in (6) from its nominal position to a new position 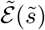 (note that the set-point *x*_d_ in (8) used by the algorithm depends on 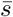). Hence, the solutions of the closed-loop system will converge to some other point 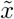 to which it corresponds a different ratio at steady state between the species 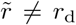. Therefore, we can analyze the robustness of the controllers by evaluating the relationship between the steady-state ratio 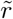 and the perturbed 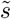.

To analyze the robustness of the state-feedback controller (9) we sampled the parameter space 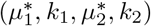 with the orthogonal sampling method [15] by varying each parameter by ±30% from its nominal value and taking 1000 samples. For each sample 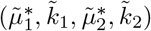 we evaluated the corresponding 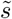 and the ratio 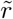 to which the closed-loop system converge. The result of this numerical analysis is reported in Fig. 5, left panel. We can notice that perturbations in the parameters that cause an error on 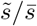 at most of 50% result in a regulation error 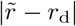 smaller than 30%.

**Fig. 5.**
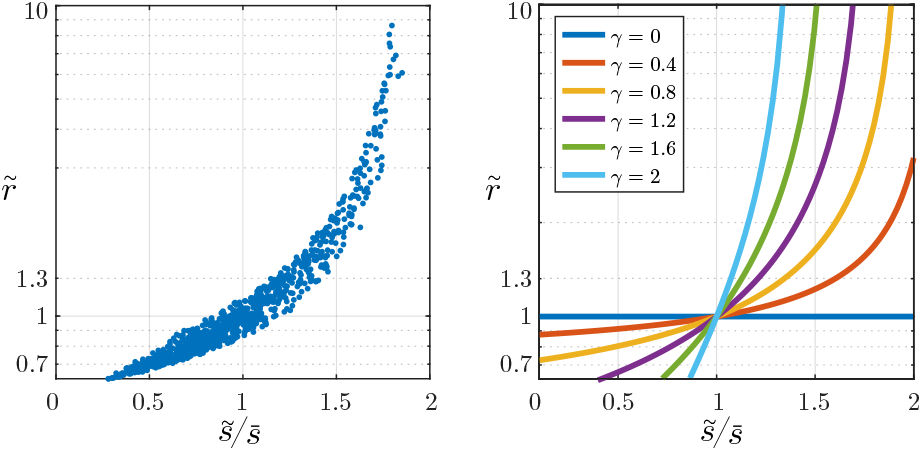
Left panel: Robustness analysis of the state-feedback controller. For each sample 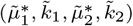 in the parameter space, the substrate concentration at coexistence 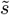 and the steady-state ratio 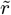 in perturbed conditions were evaluate at closed-loop. A total of 1000 samples were extracted using orthogonal sampling. Right panel: Robustness analysis of the switching controller. The graph of the function 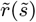 is reported for different values of the control parameter *γ*. The control parameters of the algorithms were set as in Figs. 3-4, with desired ratio *r*_d_ = 1.

For what concerns the switching controller, under perturbation of the parameters, the control algorithm is such made that it still guarantees convergence to the point of intersection between Σ and the perturbed equilibrium set 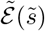. This point can be analytically computed to explicitly obtain the following relationship between 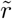 and 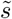, that is, 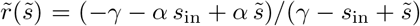. The graph of the function 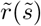 is reported in Fig. 5, right panel, for different values of the control parameter *γ*. As the control parameter *γ* decreases, the closed-loop response is more robust to perturbations of the growth functions. Moreover, it can be noticed that for *γ* ≤ 0.8 the switching controller already exhibits higher robustness that the state-feedback controller.

The performance of the control algorithms have been compared by numerically evaluating the average settling time 〈*T*〉 of their closed-loop response as their control parameters change. As the settling time strongly depends on the initial condition we choose, for each value of the control parameters, 〈*T*〉 was computed by uniformly sampling the domain 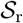, evaluating the settling time taking each point as initial condition and then averaging the results over all initial conditions. For the state-feedback controller, we evaluated 〈*T*〉 as the desired closed-loop eigenvalues (λ_1_, λ_2_) are varied over [–2,0] × [–2,0]. The results are reported in Fig. 6, left panel, showing a minimum value of about 16 h. Smaller values of the eigenvalues resulted in strongly saturated control inputs and therefore they were not considered. Analogously, the average settling time 〈*T*〉 for the switching controller was evaluated as the control parameter *γ* was varied in the interval [0.05,2]. The results are reported in Fig. 6, right panel, showing a minimum value of about 16.87 h at *γ* ≈ 0.35.

**Fig. 6.**
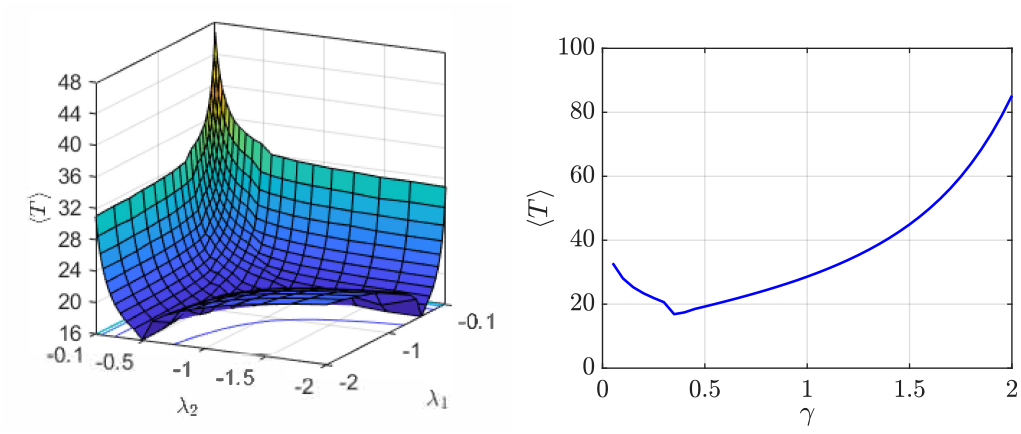
Average settling time 〈*T*〉 as function of the tuning parameters of the state-feedback control (left panel) and switching control (right panel) strategies.

## V. Conclusion

We presented the analysis and control of the concentrations of two cellular populations growing in continuous culture in a single chemostat. By exploiting the nonlinear dynamics of the open-loop system we constructed two alternative control strategies, a gain-scheduling state feedback controller and a sliding controller. We showed that both are effective in solving the problem while guaranteeing robustness to uncertainties and parameter variations. A pressing problem is that when such strategies are adopted, while the desired reference ratio between the two populations can be achieved, the convergence time is partially limited by the populations own dynamics. Also, sliding controllers are not easy to implement in this context and appropriate solutions will have to be explored to make the switching frequency compatible with the physical constraints on the control input *D*(*t*) Moreover, real-time measurements of population densities required to compute the feedback control inputs can be challenging to obtain, due to noise and cell-to-cell variability in the expression of the reporter proteins. To solve part of these problems, ongoing work is aimed at modeling and controlling multi-chamber chemostat design. For example, in a three chamber setting, each population could be grown in a separate vessel and a multi-input control strategy could be devised to control their ratio in the third chamber where they are mixed together.

